# Coral assemblages at higher latitudes favour short-term potential over long-term performance

**DOI:** 10.1101/2021.09.29.462350

**Authors:** James Cant, James D. Reimer, Brigitte Sommer, Katie M. Cook, Sun W. Kim, Carrie A. Sims, Takuma Mezaki, Cliodhna O’Flaherty, Maxime Brooks, Hamish A. Malcolm, John M. Pandolfi, Roberto Salguero-Gómez, Maria Beger

## Abstract

The persistent exposure of coral communities to more variable abiotic regimes is assumed to augment their resilience to future climatic variability. Yet, while the determinants of coral population resilience across species remain unknown, we are unable to predict the winners and losers across reef ecosystems exposed to increasingly variable conditions. Using annual surveys of 3171 coral individuals across Australia and Japan (2017-2019), we explore spatial variation across the short- and long-term dynamics of competitive, stress-tolerant, and weedy assemblages to evaluate how thermal variability mediates the structural composition of coral communities. We illustrate how, by promoting short-term potential over long-term performance, coral assemblages can reduce their vulnerability to stochastic environments. However, compared to stress-tolerant, and weedy assemblages, competitive coral taxa display a reduced capacity for elevating their short-term potential. Accordingly, future climatic shifts threaten the structural complexity of coral assemblages in variable environments, emulating the degradation expected across global tropical reefs.

## Introduction

Anticipating the resilience of natural communities requires an in-depth understanding for the determinants underpinning their constituent populations’ responses to recurrent disturbances^1,2^. Changes in environmental regimes provoke spatial shifts in the performance and distribution of populations, which upscale to the compositional reassembly of biological communities^3,4^. Exposure to more variable environments is expected to indirectly augment community resilience^5,6^. However, nuanced relationships between population characteristics and biophysical conditions ensure inconsistent responses to climate shifts^7^; with differential population sensitivities to habitat change having both accelerated and reversed expected poleward range shifts in response to climate warming^8^. By linking the mechanisms driving heterospecific variation across population responses to environmental change one can predict the resilience of whole communities to increased climatic variability^2,9,10^.

Located at the interface between tropical and temperate ecoregions, subtropical coral communities provide an opportunity for exploring the determinants of population resilience^11–13^. Recently, subtropical coral communities have undergone transformation with tropical coral taxa undergoing poleward range expansions in response to shifting thermal regimes^14–18^. At higher latitudes, however, coral communities are exposed to enhanced seasonality and cooler temperatures, and thus experience greater abiotic variability relative to their tropical counterparts^19^. Subsequently, subtropical coral communities offer insight into how differing coral assemblages utilise strategies to mediate their performance in response to environmental stochasticity across community- and regional-scales.

Exploring the performance of populations exposed to environmental stochasticity requires a consideration of their transient (*i*.*e*., short-term) dynamics^20–23^. Asymptotic (*i*.*e*., long-term) population growth rate (*λ*), which describes temporal changes in population size at stationary equilibrium^24^, is the predominant metric used to quantify population performance^24,25^. However, stochastic conditions maintain natural populations within a transient state, preventing the emergence of stationary equilibria^20,23,26^. Within stochastic environments, recurrent disturbances impose short-term changes upon the structure of populations that can elevate (*amplify*) or diminish (*attenuate*) their growth rates, resulting in population performance characteristics deviating from long-term expectations^21,27^. Quantifying how transient population performance deviates from long-term expectations (henceforth *transient potential*) is therefore crucial for predicting the success or failure of natural populations ^28^, an approach that remains neglected within coral research^22^.

In species rich communities, evaluating ecological dynamics requires a trait-based approach to condense vast quantities of demographic detail^29^. Given the diversity of coral communities, exploring patterns across the demographic characteristics of co-occurring species presents a logistical challenge^30^. Yet, this is a challenge that can be navigated by pooling individuals based on shared trait characteristics. Morphological, physiological and phenological functional traits influence the fitness of individuals and thus determine the demographic characteristics of their populations^31^, their responses to disturbances^32^, and subsequently the assembly of biological communities^33–35^. Indeed, functional trait characteristics impact upon the demographic properties of coral populations (e.g., colony growth and reproduction^36,37^), mediating their ability to respond to local abiotic patterns^38^. Given such strong links between coral traits and demographic performance, the categorisation of coral taxa into competitive, stress tolerant, generalist and weedy life history assemblages (*sensu* ^39^) can be used to evaluate broadscale patterns in coral community reassembly^40–42^. Trait-based assessments of coral community assembly also better inform upon the wider implications of ongoing community shifts than taxonomic-based assessments, thereby aiding the management of coral reef ecosystems^42^.

Here, we investigate how the performance characteristics of tropical and subtropical coral populations map onto patterns of thermal variability across assemblages of competitive, stress-tolerant, and weedy taxa. Using Integral Projection Models (IPMs)^43^, we quantify the association between different dimensions of thermal variability (monthly mean sea surface temperature [SST], monthly SST variance, and monthly SST frequency spectrum) and the transient potential and long-term performance characteristics of tropical and subtropical coral assemblages across both the northern and southern hemispheres (Fig. 1). Specifically, we anticipate that, compared to their tropical counterparts, subtropical coral assemblages will prioritise transient potential over long-term performance, corresponding to the need for subtropical coral populations to endure periodically disturbed environments. Thus, we expect that characteristics associated with amplification capacity and transient potential will align with cooler, more variable environments, whilst measures of long-term performance will be greater in warmer, more consistent environments; a pattern that will persist irrespective of functional strategy and geographic location.

**Figure 1.**
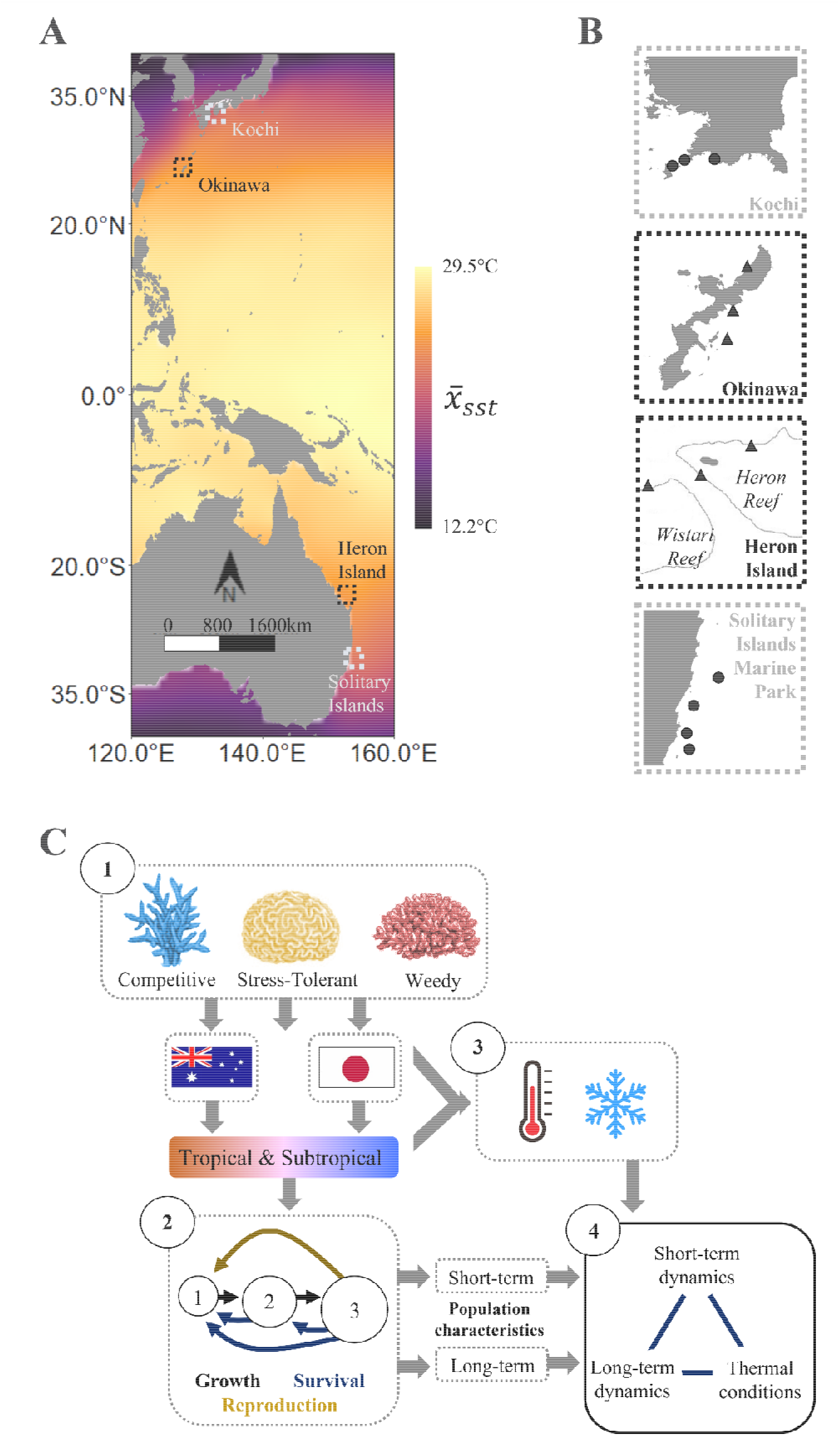
Using repeated annual surveys of tagged individual colonies, conducted between 2016 and 2019, we quantified the influence of environmental stochasticity on the long-term performance and transient potential of tropical and subtropical coral populations in southern Japan and eastern Australia. **(A)** As climate shifts induce range expansions in many coral species worldwide, their populations are increasingly exposed to a gradient in thermal regimes, illustrated here by mean monthly sea surface temperatures (x□_sst_; °C) recorded between 1950 and 2019^91^. **(B)** Between 2016 and 2019, we documented the survival, growth, fragmentation, and recruitment patterns of 3171 tagged coral individuals within the tropical reef communities (▴) of Okinawa (Japan) and Heron Island (Australia), and within the subtropical coral communities (•) of Kochi (Japan) and the Solitary Islands Marine Park (Australia). **(C)** Using this data, we parameterised Integral Projection Models (IPMs) describing the dynamics of tropical and subtropical assemblages of competitive, stress-tolerant, and weedy coral taxa. Combining outputs obtained from these models with measures of the thermal regimes experienced by each population we then explored the relationships between the long-term performance and transient (short-term) potential of coral populations, and their thermal exposure.

## Results and Discussion

Our analyses reveal contrasting patterns in long-term performance and transient potential corresponding with the exposure of coral populations to thermal variability along a gradient from warmer, more stable environments to cooler, more variable conditions (Fig. 2). Using partial least squares regression (PLSR), we evaluated how patterns in the long-term performance, demographic recovery, and transient potential, of coral populations conform with their exposure to thermal variability. We obtained estimates of long-term population performance (asymptotic population growth rate, *λ*), demographic recovery (damping ratio [*ρ*], *i*.*e*., a relative measure of the time needed for a population to converge to a stable equilibrium^24^, and transient potential (transient envelope [*TE*], *i*.*e*., the difference between maximum and minimum population size following disturbance^27,45^) from IPMs depicting the dynamics of tropical and subtropical assemblages of competitive, stress-tolerant, and weedy coral taxa in Japan and Australia (Fig. 1; Supplementary S1 & S2). Meanwhile, we quantified the exposure of these assemblages to thermal variability using three measures of local SST regimes: monthly mean SST (*x*□_*sst*_), monthly SST variance (*cv*_*sst*_), and monthly SST frequency spectrum (*β*_*sst*_; Supplementary S3).

**Figure 2.**
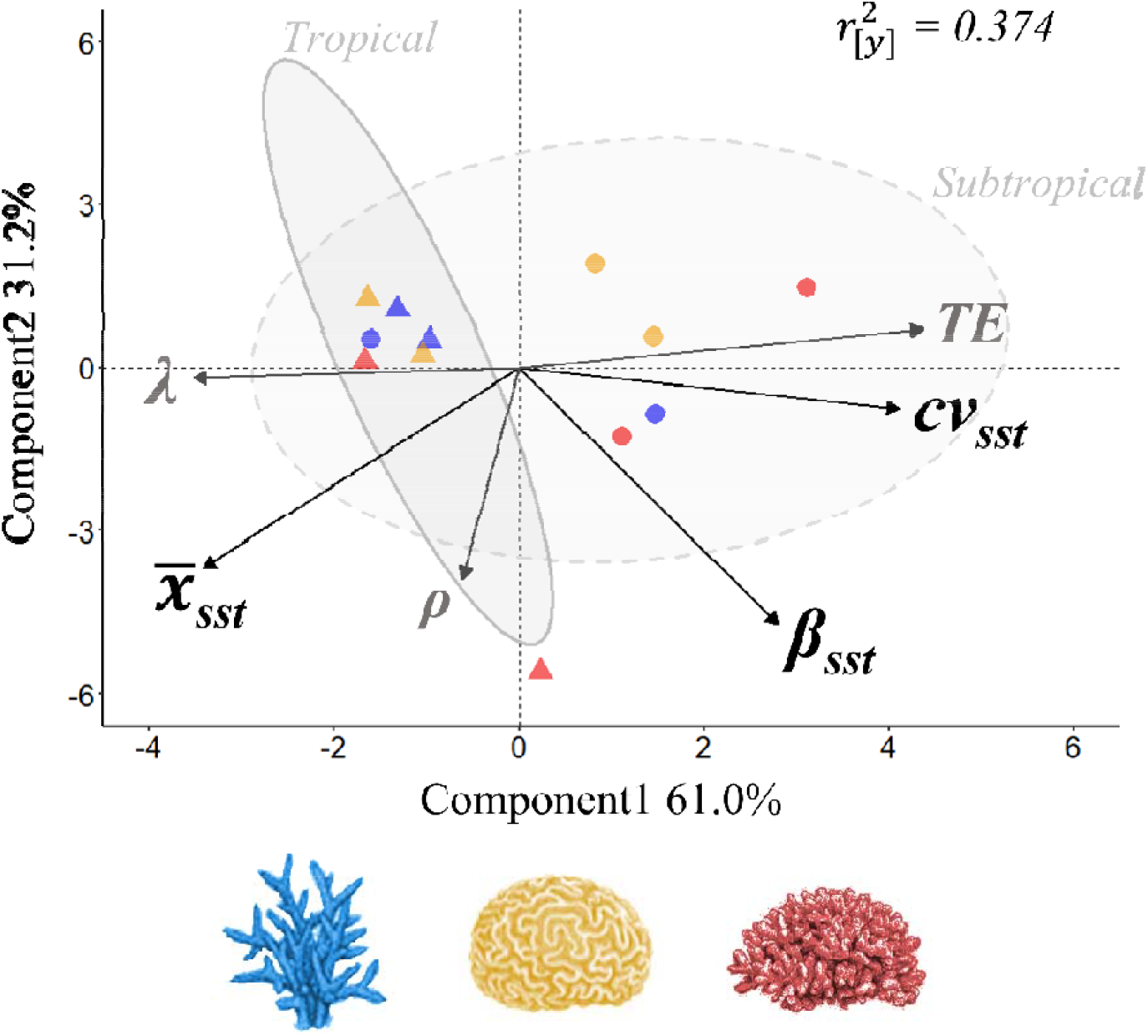
Contrasting patterns in long-term performance and transient potential across our examined coral populations, corresponding with their relative exposure to thermal variability. Partial least squares regression **s**core plot illustrating the association between thermal conditions, and the long-term performance (*λ*) and transient potential (transient envelope [*TE*] & damping ratio [*ρ*]) of tropical (▴) and subtropical (•) populations of competitive (blue), stress-tolerant (yellow), and weedy (red) coral taxa. To quantify the thermal conditions experienced by each coral population, we used sea surface temperatures (SST) recorded between 1950 and 2019 to calculate regional estimates of mean monthly SST (*x*□_*sst*_), monthly SST variance (*cv*_*sst*_), and monthly SST frequency spectrum (*β*_*sst*_). Component scores illustrate the relative degree of variance explained in the thermal predictor variables, whilst 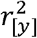 reflects the cumulative variance explained across the demographic characteristics. The shaded polygons reflect the clustering of tropical and subtropical populations, whilst the dotted lines delineate regions of association to facilitate the visualisation of patterns in correlation between the abiotic and demographic variables.

Notably, we illustrate how coral assemblages exposed to more variable thermal conditions display enhanced transient potential. Using data focused on a single taxon, Cant et al. ^44^ suggested that a capacity for short-term increases in population growth observed in a subtropical *Acropora* spp. assemblage may underpin its viability in more variable high-latitude environments. Here, we present evidence that this compensatory strategy is not only isolated in competitive coral taxa and does indeed correspond with the exposure of subtropical coral assemblages to more variable environments. Explaining 92.17% of the variance across our three measures of thermal exposure (*x*□_*sst*_, *cv*_*sst*_, and *β*_*sst*_), our PLSR captures 37.43% of the variance in long-term performance (*λ*), demographic recovery (*ρ*), and transient potential (*TE*; Fig. 2, 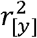). The first PLSR component depicts a gradient in SST variability, describing 60.97% of the variance in thermal conditions experienced by our examined coral assemblages. It is along this component that divergent patterns within estimates of *λ* and *TE* are most pronounced. Consequently, estimates of *TE* are positively correlated with the measures of thermal variability (*cv*_*sst*_) and frequency spectrum (*β*_*sst*_), whilst higher *λ* estimates associate with warmer mean monthly SSTs (*x*□_*sst*_; Fig. 2). Meanwhile, damping ratio (*ρ*) estimates are aligned with the second PLSR component that describes patterns in mean SST (*x*□_*sst*_) and SST frequency (*β*_*sst*_). Enhanced transient potential is thought to buffer the performance of populations exposed to elevated abiotic variability, elevating their capacity to exploit more variable environments^46,47^. However, variation in transient potential across our assemblages of differing coral taxa (Fig. 3), suggests that exposure to abiotic variability alone does not assure resilience towards future climatic variability.

**Figure 3.**
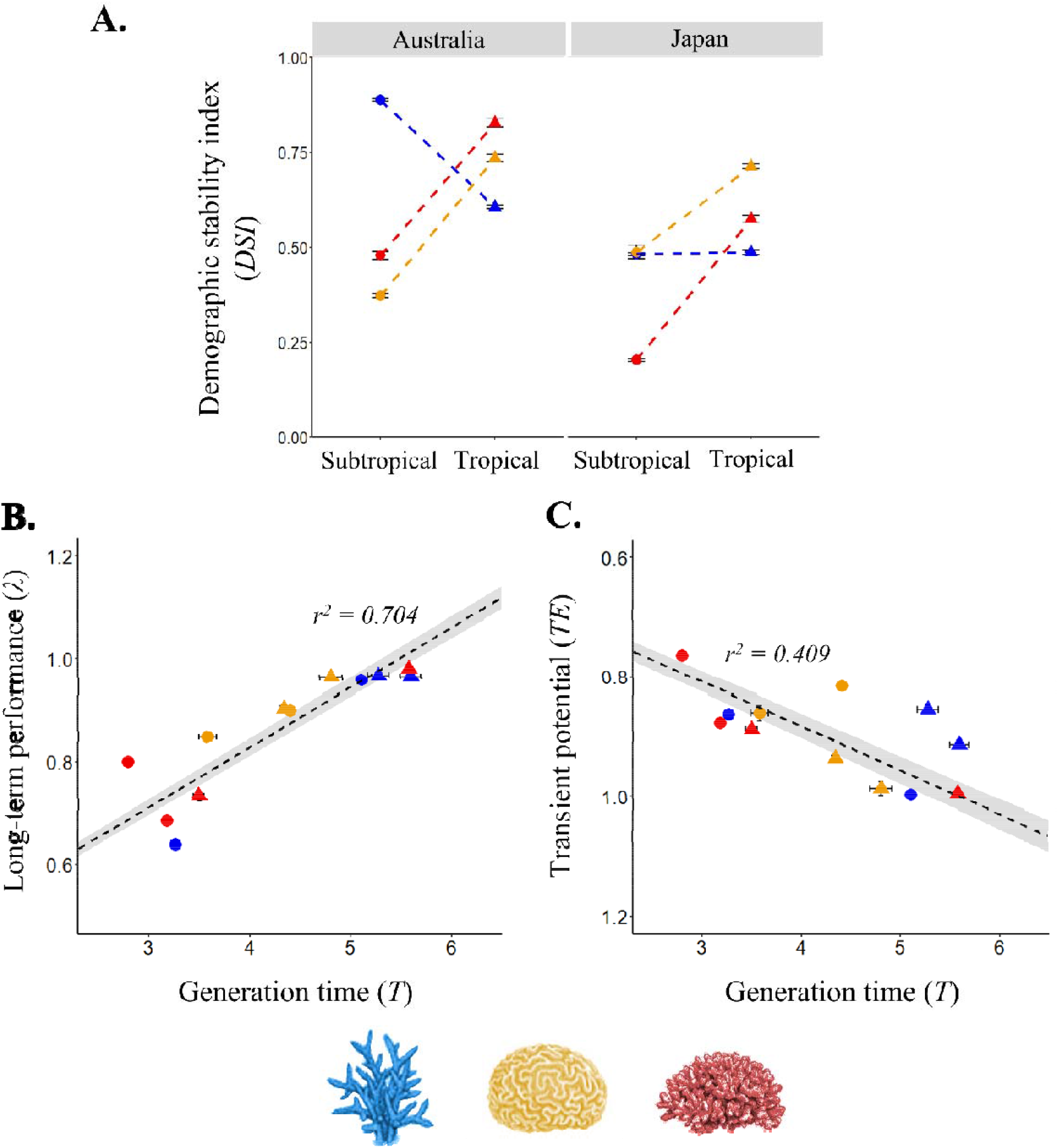
Inter-specific variation within the contrasting patterns observed between long-term performance and transient potential across tropical and subtropical correlates with patterns in population turnover rate. **(A)** Interaction plot showcasing how estimates of demographic stability index (DSI) vary between associated tropical (▴) and subtropical (•) populations of competitive (blue), stress-tolerant (yellow), and weedy (red) coral taxa in Australia and Japan. We present DSI, as an inverse measure of maximal amplification (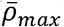), describing the ability for populations to undergo elevated growth following disturbance. Thus, lower DSI estimates correspond with enhanced amplification capacity. We also applied Type 2 linear regression to separately explore the association of population turnover characteristics with **(B)** long-term performance (asymptotic population growth rate; *λ*), and **(C)** transient potential (transient envelope, *TE*) across tropical and subtropical populations of competitive, stress-tolerant, and weedy coral taxa in Australia and Japan. We note here that transient envelope estimates were reversed during transformation to achieve normality, thus higher values reflected diminished transient potential. We have therefore displayed transient potential on a reversed scale to facilitate comparisons with patterns in long-term performance (*λ*). We used generation time (Years; displayed here on the log scale) as a measure of population turnover rate, with higher estimates reflecting slower rates of population turnover. Across panels B and C *r*^*2*^ values are provided as measure of model fit. Across all panels error is displayed using 95% CI.

Although evident across taxa, the contrasting pattern we observe between long-term performance and transient potential does not manifest consistently between paired tropical and subtropical coral assemblages (Fig. 3A & Table 1). We explored inter-assemblage variation across estimates of long-term performance and transient potential. Again, we quantified long-term performance using *λ*, whilst a *demographic stability index* (*DSI*) calculated from our IPMs provided a measure of transient potential. A three-way ANOVA revealed significant interactions between the factors of assemblage classification (competitive, stress-tolerant, or weedy), ecoregion (tropical *vs*. subtropical), and country (Australia vs. Japan; ANOVA_*λ*_: F_2,11562_ = 5698.47, *p <* 0.001; ANOVA_DSI_: F_2,11581_ = 589.8, *p <* 0.001). Despite this, the tropical assemblages routinely possess higher estimates of *λ* relative to their subtropical counterparts (Tukey: *p <* 0.001 in all cases; Table 1). The one exception were weedy corals in Japan, where *λ* is highest in the subtropics (*λ*_[t]_ = 0.760 [95% CI: 0.750, 0. 770], *λ*_[s]_ = 0.807 [0.802, 0.812]; *p <* 0.001). Alternatively, our subtropical coral assemblages typically possess a greater capacity for amplifying population growth following a disturbance than our tropical assemblages (Fig. 3A). Yet, this pattern is not consistent across life history strategies, with competitive assemblages exhibiting the opposite trend in Australia (*p <* 0.001) and no variation in Japan (*p =* 0.999).

**Table 1.** Population growth rates (*λ*) obtained from corresponding tropical and subtropical populations of competitive, stress-tolerant, and weedy coral taxa in Australia and Japan. Shading used to highlight the highest estimate of population growth across each tropical-subtropical pairing. Error displayed using 95% CI.

| Country | Life-history group | Tropical | Subtropical |
| --- | --- | --- | --- |
| Australia | Competitive | 0.983 [0.981, 0.984] | 0.958 [0.957, 0.959] |
|  | Stress-tolerant | 0.983 [0.980, 0.985] | 0.899 [0.898, 0.899] |
|  | Weedy | 0.981 [0.980, 0.982] | 0.686 [0.684, 0.687] |
| Japan | Competitive | 1.001 [0.999, 1.004] | 0.640 [0.639, 0.641] |
|  | Stress-tolerant | 0.913 [0.909, 0.917] | 0.885 [0.877, 0.894] |
|  | Weedy | 0.760 [0.750, 0.770] | 0.807 [0.802, 0.812] |

The relative long-term performance and transient potential of our coral assemblages corresponds with patterns in their generation time (Fig. 3B & C). We used Type 2 linear regression^48^ to explore the relationship between estimates of generation time (*T, i*.*e*., the time needed for individuals of a population to be replaced^49^, long-term performance (*λ*), and transient potential (*TE*) calculated from our IPMs. Generation time is a strong predictor of long-term population growth rate (*r*^*2*^ _=_ 0.704), with long-term performance increasing with generation time (Fig. 3B). Conversely, longer generation times are associated with reduced transient potential (Fig. 3C; *r*^*2*^ _=_ 0.409). Hence, shifts between long-term performance and transient potential, in response to thermal variability, manifests differently across our examined tropical and subtropical coral assemblages, due to variation in their characteristics of temporal population turnover.

### Transient buffering in variable environments

Principally, contrasting patterns between long-term performance and transient potential imply that long-term performance does not predict the capacity for populations to endure repeated disturbances. Also, whilst enhanced transient potential may enable natural populations to persist within variable environments, it comes at a cost to their long-term performance. Historically, variability in population growth rate was thought to diminish individual fitness^50^, thus hindering the persistence of populations^51^. This understanding formed the basis of the demographic buffering hypothesis, whereby populations can minimise the influence of environmental stochasticity on their long-term performance by limiting temporal variability in crucial vital rates (e.g., survival, development, and reproduction^52^). Thus, variable environments were assumed to select for populations with the ability to buffer key vital rates, thereby reducing temporal variation in performance characteristics^50,52,53^. More recently, however, enhanced transient potential has been presented as an adaptive mechanism that allows populations to exploit more stochastic environments^46^. Ellis & Crone ^47^ demonstrated how increased transient potential can buffer the effects of stochastic conditions on population growth rates, an effect that was increasingly evident in populations possessing lower *λ* estimates. Thus, it is not unexpected that coral assemblages established within variable environments would possess enhanced transient potential (Fig. 2), but the energetic cost associated with this strategy would likely inhibit their long-term performance characteristics.

Our finding that transient potential is greatest in coral assemblages displaying reduced long-term performance contrasts with previous work on mammals and plants showing a positive association between population growth rates and transient potential (e.g., ^54,55^). Faster population growth rates are assumed of populations characterised by faster individual development and high fecundity^56^, with these populations also expected to exhibit greater variability in size following disturbances^55^. While each of our surveyed assemblages are in, or close to, a state of long-term decline (*λ* < 1; Table 1), projected long-term performance was highest in the tropics, where relative transient amplification was at its lowest (Fig. 3A). Populations exhibiting longer generation times typically display reduced temporal variability in size due to higher investment in individual survival reducing the need to counteract disturbances^54^; a pattern that we show to be evident in our examined coral assemblages (Fig. 3C).

### Interspecific variation in transient characteristics

Here we show that stress-tolerant and weedy coral taxa possess more pronounced transient amplification at higher latitudes, highlighting a potential mechanism supporting their persistence. Short-term increases in population growth following disturbance have been demonstrated in a representative subtropical competitive coral assemblage (*Acropora* spp.)^44^. However, subtropical-tropical variation in the amplification capacity of competitive coral assemblages appears minimal in comparison to the variation observed here across stress-tolerant and weedy assemblages (Fig. 3A). Weedy corals typically exhibit smaller colony sizes, faster growth rates, and brooding reproductive strategies, producing larvae that settle quickly after release^39,57^. Together, these strategies support faster population turnover, enabling weedy coral species to proliferate within highly disturbed environments^58^. Conversely, stress-tolerant corals display slower growth rates, longer life expectancies, high fecundity, and broadcast spawning strategies^39,59^. The larger, more robust, morphologies associated with stress-tolerant coral taxa maximise energy storage, promoting their persistence within challenging environments^60^. Longer lifespans and elevated fecundity allow stress-tolerant corals to endure stochastic conditions by taking advantage of sporadic improvements in local conditions^39^. Consequently, our findings support existing projections that weedy and stress-tolerant coral taxa are likely to become increasingly prevalent throughout disturbed coral communities^61,62^. However, these projections herald the future loss of the structural complexity considered essential to the functioning of reef ecosystems^63^.

Crucially, our findings do not reflect the current reality for many coral assemblages within regions of high abiotic variability, suggesting that the composition of coral communities is not solely mediated by the interplay between transient dynamics and abiotic variability. Despite the lower amplificatory capacity of subtropical competitive corals compared to subtropical weedy and stress-tolerant assemblages, competitive coral taxa dominate many subtropical coral assemblages^64–66^. Utilising fast growth strategies, colonies of competitive coral taxa are capable of rapidly colonising available substrate, quickly outcompeting heterospecifics for both space and light^39^. Whilst this competitive nature explains their dominance across contemporary subtropical communities, the sensitivity of many competitive coral taxa to environmental shifts means that these assemblages are often regarded as early successional, dominating only within optimal environments, and receding as reef ecosystems approach climax states^67,68^. Within subtropical environments coral community composition is mediated by environmental pressures and dispersal barriers that filter the occurrence of species according to their trait characteristics^38,69^. As a result, subtropical coral assemblages typically consist of a subset of tropical species found on tropical coral reefs^38^, as well as subtropical specialists and endemics. The dominance of competitive coral taxa within subtropical coral assemblages, despite their reduced transient performance relative to other coral taxa, may therefore, imply that competitive interactions profoundly influence the performance of coral populations^70,71^. Certainly, further investigation into the influence of competitive interactions upon the transient dynamics of coral populations is needed to disentangle how coexistence between coral populations facilitates their persistence within variable environments.

### Conclusions

A limited understanding for the abiotic determinants driving the dynamics of coral assemblages inhibits our capacity to predict their future performance and, therefore, manage global coral community reassembly^72–74^. Here, we demonstrate how coral assemblages within regions of high environmental stochasticity exhibit demographic strategies associated with enhanced transient potential, but at a cost to their long-term performance. Climatic change is exposing coral communities worldwide to increased abiotic variability. Crucially, our findings here emphasize that whilst coral assemblages can adopt demographic strategies enhancing their viability when exposed to abiotic variability, the winners and losers within future, more variable environments cannot be predicted from existing measures of long-term performance. Equally, the relationship that we observed between transient potential and thermal variability was not universal across coral taxa, nor did it manifest identically across hemispheres. Subtle patterns in the association between population dynamics and their climate drivers hinder predictions of the consequences of environmental change within biological communities^75^. Nevertheless, relative to competitive coral taxa, weedy and stress-tolerant corals appear to possess a greater capacity for enduring within environments characterised by repeated abiotic disturbances. Yet, competitive coral taxa are often associated with more complex morphologies and therefore support the structural complexity critical to the wider functioning of coral associated ecosystems^63^. Accordingly, future increases in abiotic variability threaten the viability of coral associated ecosystems.

## Methods

### Modelling population dynamics

Integral Projection Models (IPMs) capture how the state composition of individuals influences the performance of populations over discrete time periods (*t* to *t+1*)^43^. Here, to quantify the long-term performance characteristics and transient (*i*.*e*., short-term) potential of coral populations, we used IPMs describing patterns in colony survival (*σ*), transitions in size (growth and shrinkage, *γ*), fragmentation probability (*κ*), fecundity (*φ*), and recruitment (□), each as a function of colony size (*z*; visible horizontal surface area, cm^2^). Specifically, our IPMs took the form

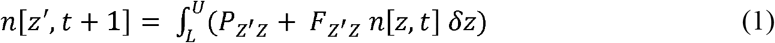

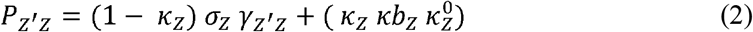

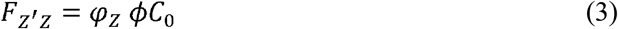

with [L, U] representing the range of possible colony sizes; calculated as 10% above and below observed maximum and minimum colony sizes to avoid accidental exclusion^76^. Accordingly, the structure of a population at time *t+1* (*n[z’, t+1]*) is a product of its structure at time *t* (*n[z’, t]*) subject to the survival (*σ*_*z*_) and transition of individual colonies from size *z* to size *z*’ (*γ*_*z’z*_); the probability of colony fragmentation (*κ*_*z*_) and the number (*κb*_*z*_) and size distribution of any colony remnants produced 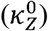; and colony fecundity (*φ*_*z*_) combined with the probability of successful recruitment (□) and the size distribution of surviving recruits (C_0_).

### Data collection

We parameterised our IPMs using data collected during repeated annual surveys of 3171 tagged colonies within tropical and subtropical coral communities in southern Japan and eastern Australia, conducted between 2016 and 2019 (Fig. 1; Supplementary S1). We tagged individual colonies using permanent plots arranged haphazardly throughout four focal coral communities (Australian subtropics [AS], Australian tropics [AT], Japanese subtropics [JS], Japanese tropics [JT]) and demarcated with numbered tags^44,62^. All tagged colonies were identified *in situ*, or from photographs, to the lowest possible taxonomic level (either genus or species). No samples were taken from tagged colonies, as although this would have allowed us to resolve species identity, we wanted to avoid any lasting interference with the processes of colony survival, growth, and fragmentation.

To facilitate comparing population characteristics observed across spatially distinct regions in Australia and Japan with varying degrees of species overlap^77^, we grouped tagged colonies across each region according to shared life-history-strategies (*sensu* ^39–41^). Specifically, we categorised colonies as ‘competitive’, ‘weedy’, ‘stress-tolerant’ or ‘generalist’ based on their morphology, growth rate and reproductive mode, following the genera classifications of Darling et al. ^39^, with minor adaptions made based on local expertise (see supplementary S2 for a detailed list). In the event that genera represented species classified across multiple categories (19 cases), we randomly assigned individuals across the relevant categories in proportion with the number of species within each category known to occur in the area (*sensu* ^41^). Following the pooling of colonies according to their life-history-strategies, we omitted all individuals defined as generalists from subsequent analyses due to their limited representation across our regional samples (n: AS = 22 colonies; AT = 31; JS = 17; JT = 65). Consequently, we constructed IPMs concerning the dynamics of each functional coral assemblage (competitive, stress-tolerant, and weedy; Fig 1) at each of the four geographical locations.

Photographs capturing the visible horizontal extent of tagged colonies were used to follow individuals over successive surveys and obtain longitudinal records of colony surface area (cm^2^; transformed to a log_10_ scale) over time. Using generalised linear mixed models (GLMMs), we estimated size-specific patterns in colony survival (*σ*), transitions in size (*γ*), and fragmentation probability (*κ*) for each population (Supplementary S1). In each case, our GLMMs included random effects (colony identity and survey location) to account for any autocorrelation between observations and within-subject variability associated with our pooling of data recorded from individuals followed across multiple years, and at different sites. Colony survival (*σ*) reflected the continued presence of tagged individuals across survey intervals (*t* to *t+1*) and was modelled as a logistic function of colony surface area at time *t*. Colony size transitions (*γ*), representing both growth through colony extension, and shrinkage through partial mortality^78^, were modelled using the polynomial relationship between initial colony surface area at time *t* and subsequent surface area at time *t+1*. Colony fragmentation probability (*κ*) was then modelled as a polynomial logistic function of colony size at time *t*. During our surveys, we recorded fragmentation in the event of observed colony breakage, recording the size (surface area, cm^2^) of all remnants produced in each case. Subsequently, we also modelled the number (*κb*_*z*_) and size 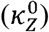 of remnant colonies produced during fragmentation as a function of colony size at time *t*, using Poisson and polynomial GLMMs, respectively.

Alongside our surveys of tagged individual colonies, we also monitored colony recruitment within our permanent coral plots. During each annual survey, we recorded the number and size of new colonies appearing within each plot. These recruitment counts enabled us to quantify annual and regional variability in recruit densities (Table S2), as well as estimate population-specific recruit size distributions (C_0_; Supplementary S1). Prior to incorporating recruitment dynamics into our IPMs, however, we first determined patterns in colony fecundity (*φ*). Using data relating colony size and larval output (larval density, cm^3^) extracted from the Coral Trait Database^79,80^, we calculated colony fecundity (*φ*) as the polynomial relationship between colony size at *t* and expected larval output (Supplementary S1). We emphasise here that this approach enabled us to complete the life cycle loop, linking the dynamics of existing individuals with the introduction of new, genetically distinct individuals within our IPMs; a necessary step when evaluating population performance^24^. However, to ensure our modelled recruitment dynamics, and therefore, our modelling framework, reflected empirical observations, we parameterised a recruit settlement function (□) into our IPMs. As a probability-based function, this recruit settlement function converts modelled larval outputs into proportional recruit densities corresponding with our empirical recruitment counts. We determined this recruit settlement function by dividing total expected larval output in any given year by the corresponding annual recruitment count (Supplementary S1, *sensu* ^62,81^).

### Quantifying population characteristics

From our IPMs, we obtained estimates of long-term performance (asymptotic population growth, *λ*), generation time (*T*), and transient potential (damping ratio [*ρ*], maximal amplification 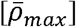 & transient envelope [*TE*]) for each tropical and subtropical coral assemblage^24,27,45,49,82^. Estimates of *λ* are typically used as a measure of long-term population viability^25^, and reflect whether a population is expected to grow (*λ* > 1) or decline (*λ* < 1) when at stationary equilibrium^24^. Generation time is a measure of population turnover, describing the time needed for individuals of a population to be replaced^49^. Alternatively, the measures of transient potential describe the expected characteristics of populations following their displacement from stationary equilibrium due to disturbances. The damping ratio constitutes a measure of demographic recovery^45,83^, describing the rate at which a population perturbed from its stationary equilibrium converges back to its asymptotic growth trajectory^24^. Meanwhile, maximal amplification quantifies the greatest increase in population size following a disturbance, relative to its asymptotic growth trajectory^27,82^. Finally, the transient envelope quantifies the magnitude by which the transient dynamics of a population deviates from its long-term trajectory^45^.

To calculate the aforementioned demographic characteristics, we discretised our IPMs into large matrices. Applying the ‘midpoint rule’ we integrated each IPM into a high-dimension matrix (200×200 cells), with the probability of transitioning from one cell to the next approximated at the cell midpoint and multiplied by the cell width as per Zuidema et al. ^84^. Estimates of *λ* were then identified as the dominant eigenvalue of each discretised matrix, whilst we estimated damping ratios as the ratio between the subdominant and dominant eigenvalues. With the *R* package *Rage*^85^ we then calculated generation time using estimates of net reproductive rate (*R*_*0*_) and *λ* obtained from each matrix,

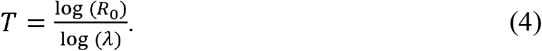

Next, we determined the transient envelope of each assemblage using their associated Kreiss bounds of amplification 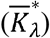 and attenuation 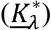,

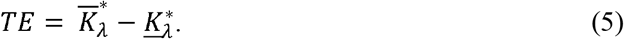

Respectively, the Kreiss bounds of amplification and attenuation reflect the largest and smallest expected long-term densities of a population following the dissipation of transient conditions, relative to its asymptotic growth trajectory^86–88^. We acknowledge here that this definition is more commonly applied to measures of population inertia^27^, which are more typically used in estimating transient envelopes^45^. However, Kreiss bound estimates have been demonstrated to align with corresponding estimates of population inertia and, unlike estimates of population inertia, are not sensitive to imprimitive population models (*i*.*e*., non-negative models permitting transitions between all state classes, but with transitions between certain stages occurring only at periodic intervals^24,27^), hence their selection here. We derived these Kreiss bounds, alongside estimates of maximal amplification, using their corresponding functions in the *R* package *popdemo*^89^.

Across each demographic measure, we determined the variance in our assemblage-specific estimates through Jack-knife resampling. During resampling, we generated 1,000 IPM variants for each assemblage, each time using 95% of our original data sample without replacement, whilst permitting recruit survival probabilities (□) to vary within observed limits. Finally, prior to their ;inclusion in further analyses, the jack-knifed distributions of *λ*, generation time, transient envelope, and maximal amplification each required transforming to ensure approximate normality. We omitted 26 variants for which *λ* > 2, as these presented unrealistic illustrations of population performance (*i*.*e*., more than doubling population size every year), before applying a log transformation to the generation time variable and a power transformation (*y*^*x*^) across the damping ratio (*y*^*-2*.*0*^), transient envelope (*y*^*-0*.*1*^) and maximal amplification variables (*y*^*-0*.*5*^).

### Evaluating spatial trends in population characteristics

To test for patterns in the spatial variation of long-term performance and transient potential across tropical and subtropical coral assemblages, we utilised partial least squares regression (PLSR), ANOVA, and Type 2 linear regression. Initially, we applied PLSR to test whether contrasting patterns in the long-term performance characteristics and transient potential of coral assemblages align with their exposure to abiotic variability. A PLSR regression quantifies the association between multiple predictor variables and one or more dependant variables^90^. Subsequently, using this technique we simultaneously evaluated the relationships between mean estimates of *λ*, damping ratio, and transient envelope obtained for each assemblage, and their correlation with patterns in thermal conditions, to provide an insight into the demographic trade-offs of coral assemblages and their mechanistic drivers.

To evaluate how abiotic variability mediates the selection of short- and long-term performance characteristics in coral assemblages, within our PLSR analyses we quantified the abiotic conditions experienced by each coral assemblage using three measures of local sea surface temperature (SST) regimes: mean monthly SST (x□_sst_), monthly SST variance (cv_sst_), and monthly SST frequency spectrum (*β*_sst_; Supplementary S3). Focusing on the four geographical regions in which our focal coral assemblages were surveyed (GPS: AS = -30.3°, 153.1°; AT = -23.4°, 151.9°; JS = 32.8°, 132.6°; JT = 26.5°, 128.1°; Fig. 1), we extracted monthly SST readings (°C; overlaid on a 1° latitude-longitude grid) taken between January 1950 and December 2019, inclusive, from the HadISST dataset^91^. Arranging these SST records into 69-year timeseries for each location, we then calculated the mean (x□_sst_) and coefficient of variance (cv_sst_) for each timeseries. Next, we estimated the frequency spectrum of each time series. Spectral analysis is used to quantify the periodicity of recurrent variability within a timeseries, with higher frequencies associated with shorter-term fluctuations^92^. The frequency spectrum of a time series is represented by its spectral exponent (*β*) and equal to the negative slope between its log spectral density and log frequency^93^, which we calculated using the package *stats*^94^. After testing these abiotic predictor variables for collinearity (Supplementary S3), we performed our PLSR analyses using the *R* package *plsdepot*^95^.

Finally, we assessed how patterns in the long-term performance, and capacity for coral assemblages to benefit from recurrent disturbance vary between tropical and subtropical regions, and how this variation manifests across coral taxa. Using a three-way ANOVA, we separately investigated variation in estimates of *λ* and maximal amplification across the three factors of country (Australia *vs*. Japan), ecoregion (tropical *vs*. subtropical), and assemblage classification (competitive, stress-tolerant, or weedy). With maximal amplification estimates inverted during transformation, larger values subsequently reflect reduced amplification potential. For the purposes of clarity in this analysis, we refer to this reversed scale as a *demographic stability index* (*DSI*), with lower values corresponding with enhanced amplification. We evaluated drivers of long- and short-term performance, by using Type 2 linear regression to separately evaluate the relationship between generation time (*T*) and estimates of *λ* and transient envelope (*TE*). Type 2 linear regression is an approach for quantifying the relationship between two non-independent variables, such that both variables include an element of error^48^. Here, due to differences in the magnitude of variance (*σ*^*2*^) across our variables of generation time, *λ*, and transient envelope (*σ*^*2*^: *T* = 1.139; *λ* = 0.009; *TE* = 0.016) we performed a Ranged Major Axis Type 2 regression using the *R* package *lmodel2*^96^.

## Acknowledgements

The authors would like to thank S. Dalton, L. Lachs, C. Kim, R. Edgar, K.-L. Gomez-Cabrera, N. Kyriacou, I. Mizukami, H. Kise, C. Fourreau, G. Masucci, P. Biondi, S. Nishihira, M. Tamae, H. Nakakoji, C. Tan, L. Lawrence, T. Hofmann, I. Montero-Serra, and all staff at Dive Quest, Marine Space, Pacific Marine, and SeaAir, for their assistance with field data collection. Additionally, aspects of this research would not have been possible without the research permits provided by the Solitary Islands Marine Park branch of the NSW Department of Primary Industries (SIMP 2016/002V2, MEAA 20/45) and the Great Barrier Reef Marine Park Authority (G19/42221.1). Funding for this research was provided by a Natural Environment Research Council (NERC) Doctoral Training Programme Scholarship to JC, a Royal Geographical Society Ralph Brown Expedition Award (RBEA 03/19) to MB and JC, the Australian Research Council Centre of Excellence for Coral Reef Studies (CE140100020) to JMP and others, the Australian Research Council Centre of Excellence for Environmental Decisions (CE110001014), a British Ecological Society small grant, the Winifred Violet Scott Charitable Trust, and the European Union’s Horizon 2020 research and innovation programme under the Marie Sklodowska-Curie grant agreement TRIM-DLV-747102 to MB. BS was supported by a Chancellor’s Postdoctoral Research Fellowship from the University of Technology Sydney and a University of Sydney Fellowship.

## Notes

### Competing Interest Statement

The authors have declared no competing interest.

### Summary of Updates

The overall manuscript has been shortened with an modification made to Figure 1 to help better clarify the methodology used.

