## Supplementary material for "Coral assemblages at higher latitudes favour short-term potential over long-term performance"

**Supplementary S1: Estimating population-specific vital rates**

Through the setup of permanent plots, we tagged colonies of competitive, stress-tolerant, and weedy coral taxa (*see below*) within the tropical and subtropical coral communities of southern Japan and eastern Australia. Across these four regional coral communities (Australian subtropics [AS], Australian tropics [AT], Japanese subtropics [JS] and Japanese tropics [JT]), permanent plots were assembled by fixing numbered tags into bare reef substrate, with each plot then consisting of a numbered tag and its surrounding coral colonies within a 2m^2^ area^1,2^. In 2016, plots were positioned throughout the subtropical Solitary Islands Marine Park (SIMP) region, Australia, at North Solitary Island (-29.93°, 153.39°), Northwest Solitary Island (-30.02°, 153.27°), South Solitary Island (-30.21°, 153.27°), Southwest Solitary Island (-30.16°, 153.23°) and Black Rock (Southwest Rocks; -30.95°, 153.08°). In 2017, further plots were then assembled in Japan, across three sites within the tropical reef communities of Okinawa (OKI; Oura Bay [26.54°, 128.08°], Hentona [26.75°, 128.18°], and Miyagi Channel [26.35°, 127.99°]) and at three sites within the subtropical communities of Kochi, Shikoku (KHI; Okinoshima [32.75°, 132.55°], Kashiwajima [32.77°, 132.62°], and Nishidomari [32.78°, 132.73°]). Finally, in 2018 plots were also arranged at three sites within the tropical reef community of Heron Island, Australia (HI; Libby’s Lair [-23.43°, 151.93°], Coral Gardens [-23.45°, 151.91°] and Wistari Reef [-23.46°, 151.87°]).

Following plot set up, annual repeated surveys of all tagged colonies, up to and including 2019, then allowed us to estimate size-specific patterns in colony survival, size transitions (growth & shrinkage^3^), fragmentation, and recruitment. Photographs, with a scale bar included for reference, were used to capture the visible horizontal extent of each tagged colony over successive surveys. With these photographs we produced longitudinal records of horizontal surface area (cm^2^) measurements for each colony using ImageJ^4^. All colony size estimates were then log-transformed to ensure a normal distribution and enhance the resolution of smaller colonies. Next, to mitigate inconsistencies in the number of census intervals across our sites in Australia and Japan we pooled data across both years and sites for each of the three

**Table S1.** Pooled number of colonies used to evaluate size-specific patterns in colony survival, transitions in size, fragmentation, and recruitment for each regional competitive, stress-tolerant, and weedy coral assemblage in Australia and Japan.

| **Life-history group** | **Country** | **Tropical** | **Subtropical** |
| --- | --- | --- | --- |
| Competitive | Australia | 207 | 217 |
|  | Japan | 103 | 446 |
| Stress-tolerant | Australia | 162 | 329 |
|  | Japan | 646 | 274 |
| Weedy | Australia | 93 | 290 |
|  | Japan | 147 | 257 |

life history categories (competitive, stress-tolerant, and weedy) at the four focal geographical locations (AS, AT, JS, and JT; Table S1). Using generalised linear mixed models (GLMMs) we then calculated size-specific patterns in colony survival, transitions in size, fragmentation probability, fecundity, and recruitment for each assemblage.

*Survival*

Colony survival represented the continued presence of tagged individuals across successive surveys. Using a binomial GLMM, we modelled the probability of colony survival as a function of colony size at time *t*, with the variables of life-history classification, country (Australia *vs.* Japan), and ecoregion (tropical *vs.* subtropical) included as fixed effects (Fig. S1). We also included the random effects of colony identity and survey location to address any within-subject-variability and autocorrelation arising from our pooling of data across multiple years and sites.

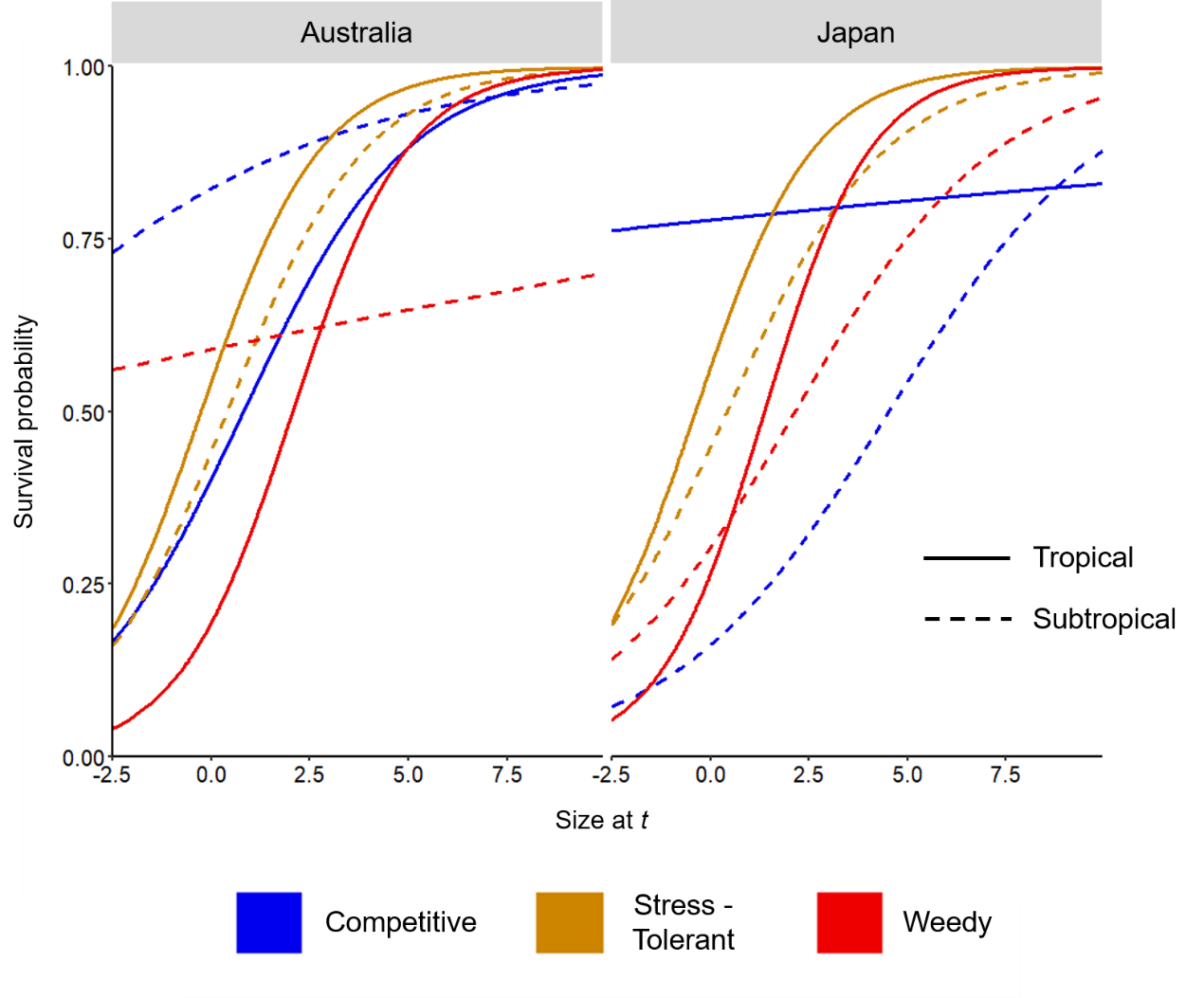

**Figure S1.** Colony survival probability as a function of colony size, showing the regional and interspecific variation in colony survival across assemblages of competitive, stress-tolerant, and weedy coral taxa in Australia and Japan.

*Size transitions*

Colony size transitions reflected the change in colony surface areas recorded across successive surveys, which we modelled as colony size at *t+1* as a function of colony size at *t* using a polynomial GLMM (Fig. S2). As with survival we modelled colony size transitions with the variables of life-history classification, country, and ecoregion included as fixed effects, and the variables of colony identity and survey location included as random effects.

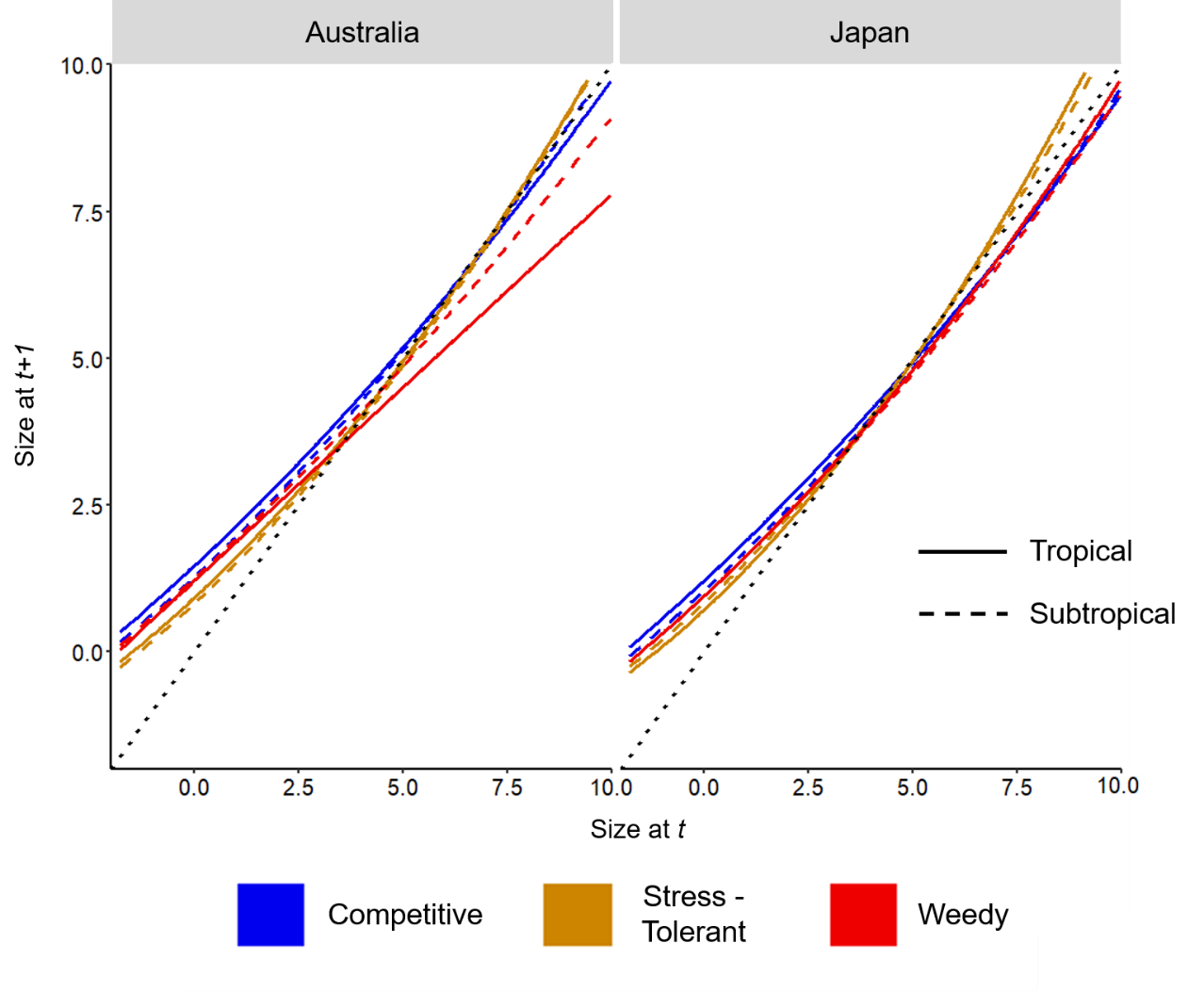

**Figure S2.** Colony size at time *t+1* as a function of colony size at time *t*, showing the regional and interspecific variation in size transition patterns across assemblages of competitive, stress-tolerant, and weedy coral taxa in Australia and Japan. 1:1 diagonal line (dotted) represents no change in size between times *t* and *t+1*.

Separately we also modelled the relationship between the variance in colony size at time *t+1* and colony size at time *t*. We determined this relationship by modelling the residuals from our initial colony size transition model as a function of colony size at time *t*, using a gamma GLMM to allow for a non-linear pattern whilst preventing negative variance. AIC scores confirmed the validity of this approach over an equivalent linear format (AIC: linear = 2413.5; gamma = 334.1). Again, we included life-history classification, country, and ecoregion as fixed effects, alongside the random effects of colony identify and site location.

*Fragmentation*

We recorded colony fragmentation in the event of observed colony breakage, recording the size (surface area, cm^2^) of all remnants produced in each case. Using a polynomial binomial GLMM, we then modelled colony fragmentation probability as a function of colony size at time *t* (Fig. S3A). Initially, we performed this analysis using only a binomial GLMM (Fig. S3B). However, despite AIC scores indicating this binomial model was the most accurate (AIC: binomial = 1167.1; polynomial binomial = 1245.1), the polynomial binomial format offered an improved representation of visual patterns within our fragmentation data (Fig. S3). As was the case across the other vital-rates, we included life-history classification, country, ecoregion, colony identity, and site location as fixed and random effects.

We also modelled the number and size of colony fragments produced during fragmentation events. With our observations of the number of fragments produced by fragmenting colonies representing count data, we modelled the number of fragments produced as a function of fragmenting colony size at time *t* using a Poisson GLMM (Fig. S4A). Meanwhile, we modelled fragment size as a function of fragmenting colony size at time *t*, using

**Figure S3.** Comparison between size-specific patterns in fragmentation probability modelled as the **(A)** polynomial binomial function of colony size at time *t*, and as the **(B)** binomial function of colony size at time *t*, showing the regional and interspecific variation across assemblages of competitive, stress-tolerant, and weedy coral taxa in Australia and Japan. Data points shown on panel A to help visualise the improved representation of observed
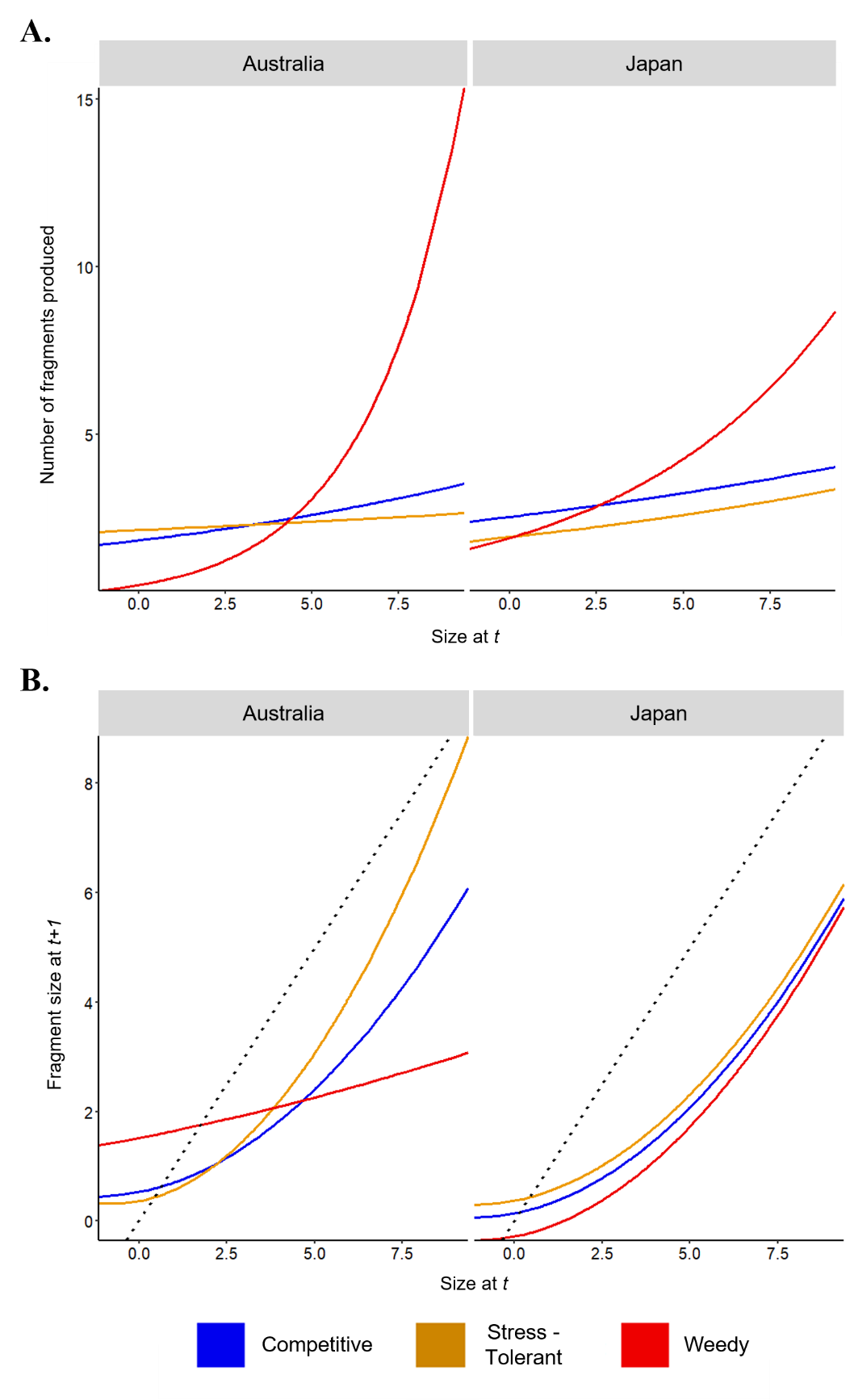
fragmentation patterns presented by the polynomial binomial model.

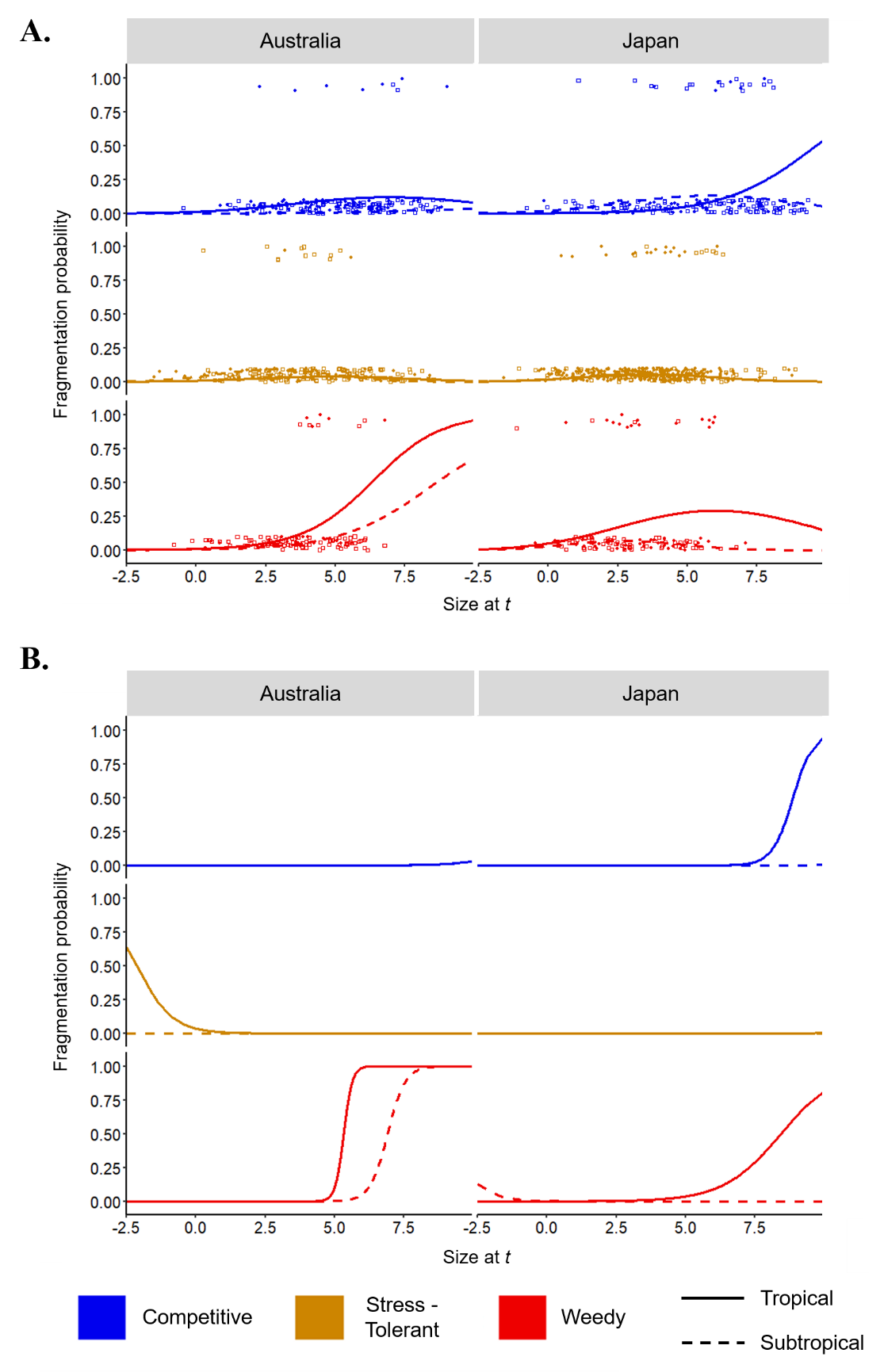

**Figure S4.** (A) Number and (B) size of fragments produced as a function of fragmenting colony size at time *t*, showing the regional and interspecific variation across assemblages of competitive, stress-tolerant, and weedy coral taxa in Australia and Japan. 1:1 diagonal line (dotted) on panel B represents the production of fragments of equal size to parent colony.

a polynomial GLMM (Fig. S4B), which provided a more representative fit than an equivalent linear format (AIC: linear = 2243.5; polynomial = 2239.1). Finally, using a gamma GLMM we also modelled the variance in fragment sizes as a function of fragmenting colony size at time *t* (AIC: linear = 1578.0; gamma = 1440.1). Across each of our models exploring size-specific patterns in the number and size of any fragments produced during fragmentation events we only included life-history classification and country as fixed effects variables. There was insufficient replication in our data for us to include the fixed effect of ecoregion and the random effects of either survey location or colony identity. Subsequently, it was necessary for our analyses to assume that size-specific patterns in the number and size of fragments produced during fragmentation events is consistent across tropical and subtropical conspecifics.

*Recruitment*

During the repeated surveys of our permanent coral plots, we recorded the number and size of new colonies appearing within each plot. Using these recruit counts we quantified annual and regional variation in the recruit densities of competitive, stress-tolerant, and weedy coral populations (Table S2). We also used the size of observed recruits to estimate assemblage-specific recruit size distributions (Fig. S5). With the parental lineage of new recruits unknown, we modelled recruit size at time *t+1* independent to colony sizes at time *t* using a linear regression, extracting the mean recruit size and standard deviation for each assemblage. Initially, we included life-history classification, country, and ecoregion as fixed effects within this recruit size model allowing us to quantify both inter-assemblage and regional variation in the size of new recruits (Fig. S5). However, the majority of the variation between each population’s recruit size distribution was solely generated by their life-history group classification (ANOVA. F_2,1108_ = 48.8, *p <* 0.001), with the country and ecoregion variables providing only a small contribution (ANOVA. Country: F_1,1108_ = 6.4, *p* = 0.01; Ecoregion: F_1,1108_ = 0.007, *p* = 0.932; Fig. S5). Subsequently, to maximise our sample size for estimating recruit size parameters we subsequently dropped both the ecoregion and country terms from the model.

**Table S1.** Densities of new colonies of competitive, stress-tolerant, and weedy coral taxa observed during the repeated surveys of our permanent coral plots in 2017, 2018, and 2019.

| **Country** | **Ecoregion** | **Population classification** | **2017** | **2018** | **2019** |
| --- | --- | --- | --- | --- | --- |
| Australia | Tropical | Competitive | - | - | 80 |
|  |  | Stress-Tolerant | - | - | 83 |
|  |  | Weedy | - | - | 59 |
|  | Subtropical | Competitive | 4 | 16 | 33 |
|  |  | Stress-Tolerant | 1 | 31 | 74 |
|  |  | Weedy | 6 | 52 | 80 |
| Japan | Tropical | Competitive | - | 12 | 31 |
|  |  | Stress-Tolerant | - | 108 | 188 |
|  |  | Weedy | - | 10 | 12 |
|  | Subtropical | Competitive | - | 34 | 73 |
|  |  | Stress-Tolerant | - | 21 | 26 |
|  |  | Weedy | - | 42 | 44 |

**Figure S5.** Regional and interspecific variation in the recruit size densities of competitive, stress-tolerant, and weedy coral assemblages in Australia and Japan.

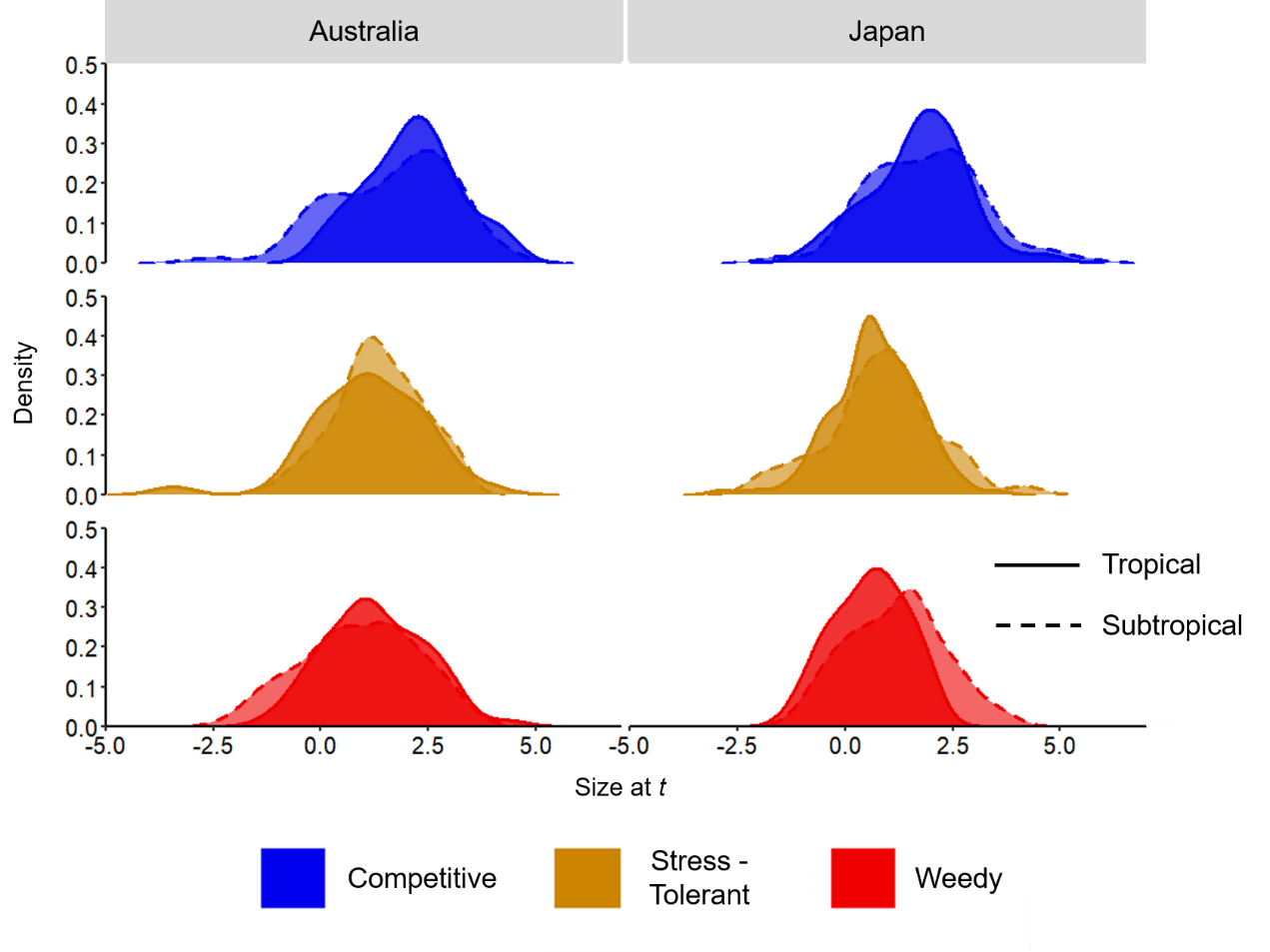

*Fecundity*

Due to the logistical challenges associated with insitu measurements of colony fecundity^5^, we did not empirically record the fecundity of our tagged colonies. Instead, we modelled size-specific patterns in colony fecundity using a relationship linking colony size and larval output (larval volume, mm^3^) recorded in the coral communities at Lizard Island, on the Great Barrier Reef^6^. Firstly, we categorised the coral species surveyed by Hall & Hughes ^6^ as competitive, stress-tolerant, or weedy according to their shared life-history characteristics (*sensu* ^7^). Using a polynomial GLMM we subsequently quantified a relationship between colony size and larval output for competitive, stress-tolerant, and weedy coral taxa (Fig. S6).

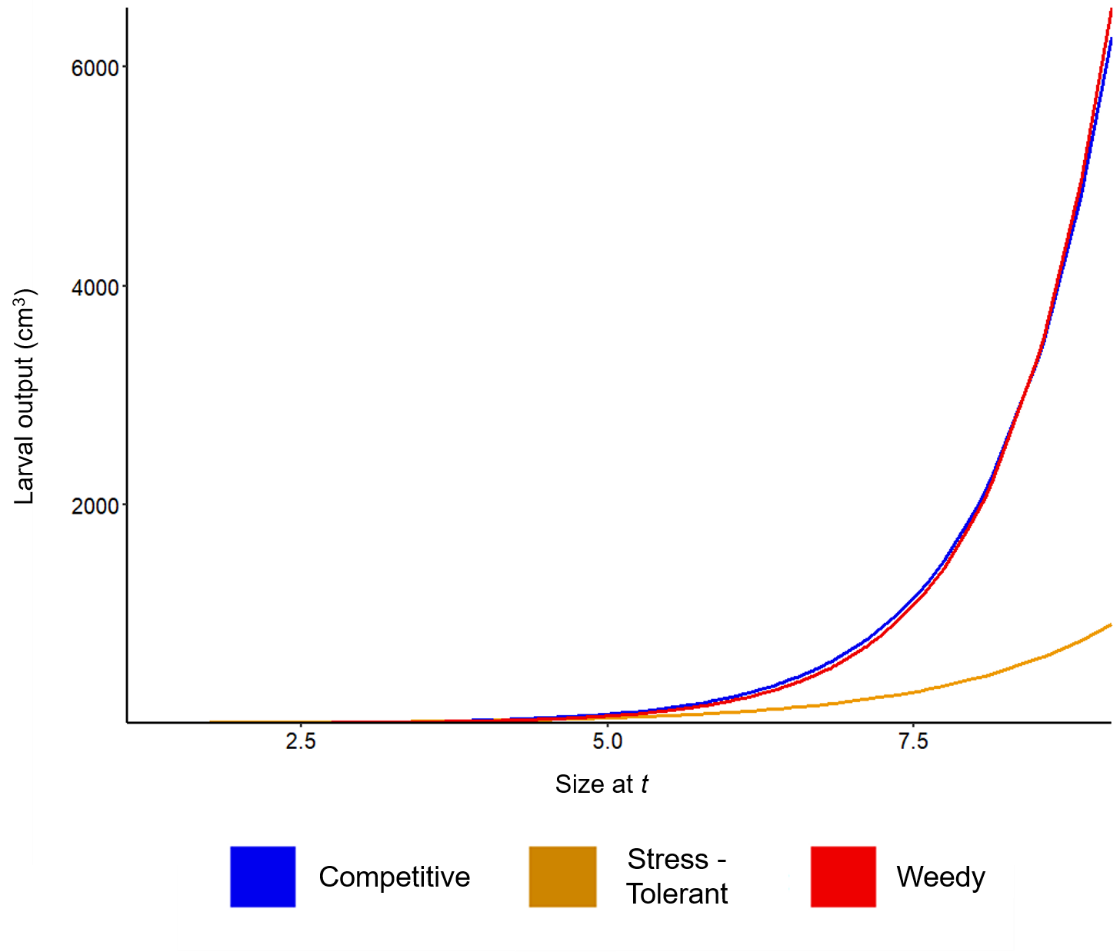

**Figure S6.** Size-specific patterns in larval output (cm^3^) estimated for competitive, stress-tolerant, and weedy coral populations using data obtained from coral communities on Lizard Island, on the northern Great Barrier Reef^6^.

We acknowledge here that our approach to modelling fecundity does imply an assumption that all larvae produced by an assemblage will reseed back into that same assemblage; an assumption that is inappropriate for coral populations which typically exist as open populations with larvae capable of dispersing away from their source populations^8,9^. We corrected this assumption by parameterising a recruit survival function (*ϕ*) into our IPMs. This recruit survival function serves to convert estimates of larval output from a measure of volume into the proportional contribution of colonies towards observed recruit densities, as a function of their size. Thus, although we have modelled fecundity using data from a distinctly different community, our use of the recruit survival function ensures that recruitment patterns within our IPMs were determined by empirical counts made within our focal communities and made no assumptions regarding the initial source of new recruits. Our IPMs were therefore not sensitive to changes in colony fecundity, and our inclusion of this vital rate merely allowed us to close the loop between the dynamics of existing colonies and the dynamics of recruitment in order to quantify measures of long-term population performance and transient potential. We estimated the recruit survival function as the ratio between the total expected larval output of a population in any given year and the corresponding annual recruitment count for that population (*sensu* ^1,10^).

**Supplementary S2: Classifying tagged corals according to shared morphological and ecological trait characteristics**

**Table S2.** Table of coral genus and species to which our tagged colonies were identified, alongside the proportion of each taxonomic rank assigned to each of the four assemblage classifications: Competitive, Generalist, Stress-tolerant, and Weedy. Greyscale used to differentiate between colonies tagged in subtropical Australia, tropical Australia, subtropical Japan, and tropical Japan.

|  | **Coral genus/species** | | **Competitive** | **Generalist** | **Stress tolerant** | **Weedy** | **Unassigned** | **Citation** |
| --- | --- | --- | --- | --- | --- | --- | --- | --- |
|  | *Acanthastrea* | *echinata* |  |  | 1.00 |  |  | 2,3 |
|  | *Acanthastrea* |  |  |  | 1.00 |  |  | 2,3 |
|  | *Acropora* | *anthoceris* | 1.00 |  |  |  |  | 2,3 |
|  | *Acropora* | *cytherea* | 1.00 |  |  |  |  | 1,2 |
|  | *Acropora* | *glauca* | 1.00 |  |  |  |  | 1,2 |
|  | *Acropora* | *hyacinthus* | 1.00 |  |  |  |  | 1 |
|  | *Acropora* | *loripes* | 1.00 |  |  |  |  | 1,2 |
|  | *Acropora* | *solitaryensis* | 1.00 |  |  |  |  | 1,2 |
|  | *Acropora* |  | 1.00 |  |  |  |  | 1,2 |
|  | *Acropora* | *valida* | 1.00 |  |  |  |  | 1,2 |
|  | *Astrea* | *curta* |  |  | 1.00 |  |  | 1 |
|  | *Cyphastrea* |  |  |  | 1.00 |  |  | 1,6 |
|  | *Dipsastraea* | *speciosa* |  |  | 1.00 |  |  | 2,3 |
|  | *Dipsastraea* |  |  |  | 1.00 |  |  | 2,3 |
|  | *Goniopora* | *djiboutiensis* |  |  | 1.00 |  |  | 2,3 |
|  | *Goniopora* | *lobata* |  |  | 1.00 |  |  | 2,3 |
|  | *Goniopora* | *norfolkensis* |  |  | 1.00 |  |  | 2,3 |
|  | *Micromussa* | *amakusensis* |  |  | 1.00 |  |  | 4 |
|  | *Micromussa* | *lordhowensis* |  |  | 1.00 |  |  | 2,3 |
|  | *Micromussa* |  |  |  | 1.00 |  |  | 4 |
|  | *Montipora* |  | 0.43 | 0.14 | 0.43 |  |  | 1,2,6 |
|  | *Montipora* | *venosa* |  |  | 1.00 |  |  | 1 |
|  | *Paragoniastrea* | *australiensis* |  |  | 1.00 |  |  | 1 |
|  | *Plesiastrea* |  |  |  | 1.00 |  |  | 2 |
|  | *Pocillopora* | *aliciae* |  |  |  | 1.00 |  | 5 |
|  | *Pocillopora* | *damicornis* |  |  |  | 1.00 |  | 1 |
|  | *Porites* | *heronensis* |  |  | 1.00 |  |  | 2,3 |
|  | *Porites* | *murrayensis* |  |  | 1.00 |  |  | 2,3 |
|  | *Porites* | *stephensoni* |  |  | 1.00 |  |  | 2,3 |
|  | *Porites* (Encrusting & Massive) |  |  |  | 1.00 |  |  | 2,3 |
|  | *Stylophora* | *pistillata* |  |  |  | 1.00 |  | 1 |
|  | *Stylophora* |  |  |  |  | 1.00 |  | 1,2 |
|  | *Turbinaria* | *frondens* |  | 1.00 |  |  |  | 1 |
|  | *Turbinaria* | *mesenterina* | 1.00 |  |  |  |  | 1 |
|  | *Turbinaria* | *patula* |  | 1.00 |  |  |  | 1,3,6 |
|  | *Turbinaria* | *radicalis* |  |  | 1.00 |  |  | 1,3,6 |
|  | *Acanthastrea* |  |  |  | 1.00 |  |  | 2,3 |
|  | *Acropora* |  | 1.00 |  |  |  |  | 1,2,3 |
|  | *Astrea* | *curta* |  |  | 1.00 |  |  | 2,3,7 |
|  | *Astreopora* |  |  |  | 1.00 |  |  | 1,2,3 |
|  | *Cyphastrea* |  |  |  | 1.00 |  |  | 1,2,3,6 |
|  | *Dipsastraea* |  |  |  | 1.00 |  |  | 2,3 |
|  | *Echinophyllia* |  |  |  | 1.00 |  |  | 1,2,3 |
|  | *Echinopora* |  |  | 1.00 |  |  |  | 1,2,3,6 |
|  | *Favites* |  |  |  | 1.00 |  |  | 2,3 |
|  | *Fungia* |  |  |  | 1.00 |  |  | 2,3 |
|  | *Galaxea* |  |  |  | 1.00 |  |  | 2,3 |
|  | *Goniastrea* |  |  |  | 0.86 | 0.14 |  | 1,2,3,6 |
|  | *Goniopora* |  |  |  | 1.00 |  |  | 2,3 |
|  | *Hydnophora* |  | 0.33 | 0.33 | 0.33 |  |  | 1,2,3,6 |
|  | *Isopora* |  |  | 1.00 |  |  |  | 3 |
|  | *Leptastrea* |  |  |  | 0.60 | 0.40 |  | 1,2,3,6 |
|  | *Leptoria* |  |  |  | 1.00 |  |  | 1,2,3 |
|  | *Lobophyllia* |  |  |  | 1.00 |  |  | 1,2,3 |
|  | *Merulina* |  |  | 1.00 |  |  |  | 1,2,3,6 |
|  | *Montipora* |  | 0.31 | 0.24 | 0.45 |  |  | 1,2,6 |
|  | *Mycedium* |  |  | 1.00 |  |  |  | 3 |
|  | *Oulophyllia* |  |  |  | 1.00 |  |  | 3 |
|  | *Pavona* | *varians* |  |  | 1.00 |  |  | 1,2 |
|  | *Platygyra* |  |  |  | 1.00 |  |  | 1,2,3 |
|  | *Pocillopora* | *damicornis* |  |  |  | 1.00 |  | 1 |
|  | *Pocillopora* |  | 0.60 |  |  | 0.40 |  | 1,2,3,6,9 |
|  | *Porites (Branching)* |  |  |  |  | 1.00 |  | 2,3 |
|  | *Porites (Encrusting & Massive)* |  |  |  | 1.00 |  |  | 2,3 |
|  | *Psammacora* |  |  | 0.80 |  | 0.20 |  | 1,2,3,6,8 |
|  | *Seriatopora* |  |  |  |  | 1.00 |  | 1,2,3 |
|  | *Stylophora* | *pistillata* |  |  |  | 1.00 |  | 1,2,3 |
|  | *Stylophora* |  |  |  |  | 1.00 |  | 1,2,3 |
|  | *Symphyllia* |  |  |  | 1.00 |  |  | 3 |
|  | *Turbinaria* | *frondens* |  | 1.00 |  |  |  | 1 |
|  | *Turbinaria* | *heronensis* |  | 1.00 |  |  |  | 3,6 |
|  | *Turbinaria* | *peltata* |  | 1.00 |  |  |  | 3,6 |
|  | *Turbinaria* |  | 0.14 | 0.71 | 0.14 |  |  | 1,2,3 |
|  | *Acanthastrea* |  |  |  | 1.00 |  |  | 2,3 |
|  | *Acropora* |  | 1.00 |  |  |  |  | 1,2,3 |
|  | *Astrea* | *curta* |  |  | 1.00 |  |  | 2,3,7 |
|  | *Astrea* |  |  |  | 1.00 |  |  | 2,3,7 |
|  | *Coscinarea* | *columna* |  |  | 1.00 |  |  | 2,3 |
|  | *Cyphastrea* |  |  |  | 1.00 |  |  | 1,2,3,6 |
|  | *Dipsastrea* |  |  |  | 1.00 |  |  | 2,3 |
|  | *Euphyllia* |  |  |  |  |  | 1.00 | 6 |
|  | *Favites* |  |  |  | 1.00 |  |  | 1,2,3 |
|  | *Goniastrea* |  |  |  | 0.86 | 0.14 |  | 1,2,6 |
|  | *Leptastrea* |  |  |  |  | 1.00 |  | 1,2,6 |
|  | *Leptoseris* |  |  |  | 1.00 |  |  | 3 |
|  | *Lithophyllon* |  |  |  |  |  | 1.00 | 6 |
|  | *Lobophyllia* |  |  |  | 1.00 |  |  | 1,2,3 |
|  | *Micromussa* |  |  |  | 1.00 |  |  | 2,3,6 |
|  | *Montipora* | *millepora* |  |  | 1.00 |  |  | 1 |
|  | *Montipora* |  | 0.31 | 0.38 | 0.31 |  |  | 1,2,6 |
|  | *Pavona* | *descussata* |  | 1.00 |  |  |  | 1 |
|  | *Pavona* |  |  | 0.60 | 0.40 |  |  | 2,3,6 |
|  | *Pectinia* |  |  |  |  |  | 1.00 | 3 |
|  | *Platygyra* |  |  |  | 1.00 |  |  | 1,2,3 |
|  | *Plesiastrea* |  |  |  | 1.00 |  |  | 1,2,3 |
|  | *Pocillopora* | *damicornis* |  |  |  | 1.00 |  | 1 |
|  | *Pocillopora* |  | 0.67 |  |  | 0.33 |  | 1,2,6 |
|  | *Porites (Encrusting & Massive)* |  |  |  | 1.00 |  |  | 2,3 |
|  | *Psammocora* |  |  | 1.00 |  |  |  | 1,2,3,6,8 |
|  | *Stylocoeniella* |  |  |  |  |  | 1.00 | 6 |
|  | *Stylophora* | *pistillata* |  |  |  | 1.00 |  | 1,2,3 |
|  | *Stylophora* |  |  |  |  | 1.00 |  | 1,2,3 |
|  | *Tubastraea* |  |  |  |  |  | 1.00 | 6 |
|  | *Acanthastrea* |  |  |  | 1.00 |  |  | 2,3 |
|  | *Acropora* | *humilus* | 1.00 |  |  |  |  | 1 |
|  | *Acropora* |  | 1.00 |  |  |  |  | 1,2 |
|  | *Astrea* | *annuligera* |  |  | 1.00 |  |  | 2,3,7 |
|  | *Astrea* | *curta* |  |  | 1.00 |  |  | 2,3,7 |
|  | *Astrea* |  |  |  | 1.00 |  |  | 2,3,7 |
|  | *Astreopora* |  |  |  | 1.00 |  |  | 2,3 |
|  | *Caulastrea* |  |  |  | 1.00 |  |  | 1,6 |
|  | *Cyphastrea* |  |  |  | 1.00 |  |  | 1,6 |
|  | *Diploastrea* | *heliopora* |  |  | 1.00 |  |  | 1 |
|  | *Dipsastraea* | *pallida* |  |  | 1.00 |  |  | 1 |
|  | *Dipsastraea* |  |  |  | 1.00 |  |  | 2,3 |
|  | *Echinophyllia* |  |  |  | 1.00 |  |  | 2,3 |
|  | *Echinopora* |  |  | 1.00 |  |  |  | 1,6 |
|  | *Favites* |  |  |  | 1.00 |  |  | 2,3 |
|  | *Galaxea* | *fascicularis* |  |  | 1.00 |  |  | 2 |
|  | *Galaxea* |  |  |  | 1.00 |  |  | 2,3 |
|  | *Goniastrea* |  |  |  | 0.86 | 0.14 |  | 1,2,6 |
|  | *Hydnophora* |  | 0.25 | 0.25 | 0.50 |  |  | 1,2,6 |
|  | *Leptastrea* |  |  |  | 0.60 | 0.40 |  | 1,2,6 |
|  | *Leptoria* |  |  |  | 1.00 |  |  | 2,3 |
|  | *Lithophyllon* |  |  |  |  |  | 1.00 | 6 |
|  | *Lithophyllon* | *undulatum* |  |  |  |  | 1.00 | 6 |
|  | *Lobophyllia* |  |  |  | 1.00 |  |  | 2,3 |
|  | *Merulina* |  |  | 1.00 |  |  |  | 3 |
|  | *Montastrea* |  |  |  | 1.00 |  |  | 2,3 |
|  | *Montipora* | *foliosa* |  |  | 1.00 |  |  | 2 |
|  | *Montipora* |  | 0.32 | 0.35 | 0.32 |  |  | 1,2,6 |
|  | *Oxypora* | *lacera* |  |  | 1.00 |  |  | 1,2,3,6 |
|  | *Pachyseris* |  |  | 1.00 |  |  |  | 1,3 |
|  | *Pectinia* |  |  |  |  |  | 1.00 | 3,6,7 |
|  | *Platygyra* |  |  |  | 1.00 |  |  | 2,3 |
|  | *Plesiastrea* |  |  |  | 1.00 |  |  | 2 |
|  | *Pocillopora* | *damicornis* |  |  |  | 1.00 |  | 1 |
|  | *Pocillopora* |  | 0.67 |  |  | 0.33 |  | 1,2,6 |
|  | *Porites (Branching)* |  |  |  |  | 1.00 |  | 2,3 |
|  | *Porites (Encrusting & Massive)* |  |  |  | 1.00 |  |  | 2,3 |
|  | *Psammocora* | *nierstraszi* |  | 1.00 |  |  |  | 2,3,6 |
|  | *Psammocora* |  |  | 0.67 |  | 0.33 |  | 2,3,6 |
|  | *Symphyllia* |  |  |  | 1.00 |  |  | 3 |
|  | *Turbinaria* | *irregularis* |  |  | 1.00 |  |  | 6 |
|  | *Turbinaria* |  | 0.17 | 0.67 | 0.17 |  |  | 1,2,3,6 |
| **Citation codes:** 1. Darling *et al.* ^7^, 2. Darling *et al*. ^11^, 3. Zinke *et al*. ^12^,  4. Ng *et al.* ^13^, 5. Schmidt-Roach *et al.* ^14^, 6. Veron *et al*. ^15^,  7. Huang *et al*. ^16^, 8. Benzoni *et al*. ^17^, 9. Schmidt-Roach *et al.* ^18^. | | | | | | | | |

**Supplementary S3: Quantifying exposure to thermal variability**

To evaluate how the long-term performance characteristics and transient potential of coral assemblages correspond with gradients in their exposure to thermal variability we calculated four measures describing the local sea surface temperature (SST) regimes experienced by each population prior to, and during, our survey period. Specifically, we focused on the four measures of mean monthly SST (x̄_sst_), monthly SST variance (cv_sst_), monthly SST autocorrelation (a_sst_), and monthly SST frequency spectrum (*β*_sst_). Using the NOAA Coastwatch ERDDAP data server (<https://coastwatch.pfeg.noaa.gov/erddap/index.html>) we sourced high resolution SST records (°C; overlaid on a 1° latitude-longitude grid), from the Met Office Hadley Centre climate dataset (HadISST)^19^. From this dataset we then extracted monthly SST readings taken between January 1950 and December 2019, inclusive, at each of the four geographical regions in which our focal coral assemblages were surveyed (see Supplementary S1; GPS: SIMP = -30.3°, 153.1°; HI = -23.4°, 151.9°; KHI = 32.8°, 132.6°; OKI = 26.5°, 128.1°).

Arranging extracted monthly SST records into 69-year timeseries for each location, we then calculated the mean (x̄_sst_), variance (cv_sst_), autocorrelation (a_sst_), and frequency spectrum (*β*_sst_) for each timeseries (Table S3). We quantified the variance of each time series using its coefficient of variation which we estimated using the corresponding function in the *raster* package^20^. We then estimated the autocorrelation of each time series using the *autocorrelation* function from the *colorednoise* package^21^. Measures of autocorrelation describe the correlation between successive elements within a series, such that positive autocorrelation reflects the condition whereby the properties of any element are closely related to those preceding it^22^. Next, we estimated the frequency spectrum of each time series. The frequency spectrum of a timeseries reflects the periodicity of any recurrent variability across the series, with higher frequencies associated with shorter-term fluctuations^23^. The frequency spectrum of a time series is equal to its spectral exponent (*β*) and calculated as the negative slope between the log spectral density and log frequency of the time series^24^. We calculated the frequency spectra of each of our SST time-series using the *spectrum* function from the *stats* R package^25^.

**Table S3.** The sea surface temperature (SST) regimes experienced by coral assemblages in tropical and subtropical regions of Australia and Japan, quantified using measures of mean monthly SST (x̄_sst_), monthly SST variance (cv_sst_), monthly SST autocorrelation (a_sst_), and monthly SST frequency spectrum (*β*_sst_). Measures estimated from 69-year SST timeseries obtained from the Met Office Hadley Centre climate dataset^19^.

| **Location** | **x̄_sst_** | **cv_sst_** | **a_sst_** | ***β*_sst_** |
| --- | --- | --- | --- | --- |
| Solitary Islands  *(-30.3°, 153.1°)* | 22.77 | 9.00 | 0.86 | -1.04 |
| Heron Island  *(-23.4°, 151.9°)* | 24.72 | 9.28 | 0.86 | -1.13 |
| Kochi  *(32.8°, 132.6°)* | 22.14 | 17.10 | 0.86 | -0.94 |
| Okinawa  *(26.5°, 128.1°)* | 25.13 | 11.37 | 0.85 | -0.90 |

Finally, prior to conducting partial least squares analyses into the association between the long-term performance and transient potential of coral assemblages with patterns in thermal conditions it was necessary for us to evaluate for collinearity across our abiotic variables. We tested for collinearity using the measure of tolerance which describes an inverse measure of the correlation between multivariate predictor variables with estimates of <0.1 evidence of collinearity^26^. We calculated measures of tolerance for our abiotic variables using the function *multicol* from the *fuzzySim* package^27^. Our test for multicollinearity, when we included all four SST variables, returned tolerance estimates of ~0 highlighting a strong correlation between one or more of the variables. Subsequently, we explored collinearity across each triple-wise combination of our four abiotic variables and determined that the triple-wise combination of the variables of mean monthly SST, monthly SST variance, and monthly SST frequency spectrum exhibited the least collinearity (Table S4). Accordingly, we omitted the variable of monthly SST autocorrelation (a_sst_) from further analyses.

**Table S3.** Tolerance estimates obtained for each of the sea surface temperature (SST) measures of mean monthly SST (x̄_sst_), monthly SST variance (cv_sst_), monthly SST autocorrelation (a_sst_), and monthly SST frequency spectrum (*β*_sst_) across each triple-wise combination possible with the four abiotic variables.

| **Combination** | **x̄_sst_** | | **cv_sst_** | | **a_sst_** | ***β*_sst_** | |
| --- | --- | --- | --- | --- | --- | --- | --- |
| 1 | 0.12 | - | | 0.03 | | | 0.04 |
| 2 | - | 0.58 | | 0.22 | | | 0.17 |
| 3 | 0.26 | 0.26 | | 0.31 | | | - |
| 4 | 0.61 | 0.39 | | - | | | 0.55 |
